# Simvastatin Preserves Human Microvascular Endothelial Barrier Function Through Mevalonate-Dependent Regulation of RhoB Localization and Activation

**DOI:** 10.64898/2026.10.28.762609

**Authors:** Alejandra Morales-Maldonado, Jordan S. Pober, Richard W. Pierce

## Abstract

Endothelial barrier failure contributes to edema, tissue hypoperfusion, and organ dysfunction during inflammatory critical illness. RhoB has emerged as a regulator of tight junction disruption in human microvascular endothelial cells, but whether clinically used drugs can modulate RhoB-dependent barrier failure remains unclear. We tested whether simvastatin preserves human dermal microvascular endothelial cell barrier function during TNF stimulation and examined the role of mevalonate-dependent RhoB localization and activation. Simvastatin pretreatment attenuated TNF-induced loss of transendothelial electrical resistance and preserved junctional ZO-1 and claudin-5 organization. Simvastatin reduced membrane-associated and increased nuclear RhoB, decreased active RhoB, and suppressed myosin light chain phosphorylation. RhoB silencing phenocopied the barrier-protective effect of simvastatin, and combined RhoB silencing and simvastatin did not provide additive protection. Mevalonate supplementation reversed simvastatin-mediated preservation of barrier function, restored peripheral RhoB localization, and restored MLC phosphorylation. These findings identify mevalonate-dependent RhoB spatial regulation as a statin-sensitive mechanism contributing to inflammatory microvascular barrier dysfunction.

## INTRODUCTION

Endothelial cells (<u>**ECs**</u>) line the vascular lumen and regulate blood flow, hemostasis, immune-cell trafficking, and exchange between blood and tissues[1]. The selectively permeable barrier is maintained via inter-EC tight junctions (<u>**TJs**</u>). Endothelial barrier integrity is maintained by coordinated intercellular junctions and cytoskeletal forces that regulate paracellular permeability. In microvascular beds with prominent tight-junction (<u>**TJ**</u>) organization, proteins including ZO-1 and claudin-5 contribute importantly to this barrier. [2]. During acute inflammatory illness, TJs are disrupted and endothelial cytoskeletal tension increases, allowing inappropriate movement of plasma and proteins into the interstitial space[3]. This process, clinically recognized as capillary leak, contributes to edema, tissue hypoperfusion, and organ dysfunction in sepsis, acute respiratory distress syndrome, trauma, and many other critical illnesses[4].

Capillary barrier integrity is regulated, in large part, by Rho-family small GTPases, which coordinate actin remodeling, junctional organization, and contractile force[5]. Rho GTPases, initially described in 1985[6], belong to the Ras superfamily of small GTP-binding proteins and act as molecular switches that cycle between active GTP-bound and inactive GDP-bound states. In humans, 22 Rho family members have been identified. RhoA, Rac1, and Cdc42 have been well established as key regulators of permeability in human umbilical vein endothelial cells (<u>**HUVECs**</u>)[7]. However, HUVECs do not form high-resistance, TJ-dependent barriers like human dermal microvascular endothelial cells (<u>**HDMECs**</u>)[8]. These differences suggest that distinct Rho-family signaling mechanisms may regulate permeability in the tight-junction-forming capillary endothelium.

RhoB has emerged as a critical regulator of capillary permeability through converging human genetic, expression, and mechanistic studies. We previously identified a fatal pediatric systemic capillary leak syndrome associated with a mutation in p190BRhoGAP, a negative regulator of RhoB. This mutation prolonged RhoB activation and was associated with recurrent vascular leak, edema, hemoconcentration, hypoalbuminemia, and hypovolemic shock[9]. RhoB expression is also increased in inflamed and hyperpermeable human vascular beds, including intestinal and hepatic capillaries[10]. At the cellular level, RhoB silencing protects HDMECs from TNF-induced TJ disassembly and barrier loss[11]. Together, these findings support a model in which dysregulated RhoB activation contributes to inflammatory microvascular barrier failure.

Beyond its direct effects on endothelial barrier integrity, RhoB has also emerged as an important regulator of inflammatory signaling. Unlike other Rho family members, RhoB is rapidly induced by pro-inflammatory stimuli, including interleukin (<u>**IL**</u>)-1 and tumor necrosis factor (<u>**TNF**</u>) through nuclear factor kappa-light-chain-enhancer of activated B cells (<u>**NF-κB**</u>) activation[10], as well as by toll-like receptor ligands such as lipopolysaccharide, CpG, poly(I:C) and Rubella virus [12-14]. Interestingly, RhoB has also been implicated in the activation of the NF-κB and mitogen-activated protein kinase (<u>**MAPK**</u>) signaling pathways, leading to increased production of pro-inflammatory cytokines, including TNF, IL-1β, and IL-6[12, 13] and RhoB[11]. Unique among the Rho family, RhoB has rapid turnover and is dynamically regulated at the level of expression[15]. Through these mechanisms, RhoB activation may act as a positive feedback loop to amplify inflammatory signaling and promote capillary leak.

Given these unique roles in inflammation, RhoB represents an attractive target to block capillary leak. RhoB activity, like other Rho GTPases[16], depends on two closely interconnected regulatory processes; control of its activation state and control of its subcellular localization[10]. The activation state of Rho GTPases is governed by three classes of regulatory proteins: guanine nucleotide exchange factors (<u>**GEFs**</u>), which promote the exchange of GDP for GTP and thereby activate the GTPase; GTPase-activating proteins (<u>**GAPs**</u>), which stimulate GTP hydrolysis and return the protein to its inactive state; and guanine nucleotide dissociation inhibitors (<u>**RhoGDIs**</u>), which bind inactive GTPases and sequester them in the cytosol. In endothelial cells, only a limited number of RhoB regulatory proteins have been defined including activators ArhGEF3 [17], ArhGEF10[11] and inhibitor ArhGAP5[9]. This lack of understanding of regulation of RhoB nucleotide binding and hydrolysis make direct pharmacologic targeting challenging; modulating the mechanisms that direct RhoB localization is an attractive alternative strategy.

RhoB localization is largely directed by post-translational lipid modification of its C-terminal CAAX motif, which promotes association with plasma membrane and intracellular membrane compartments [18]. Statins are well known to inhibit β-hydroxy β-methylglutaryl-CoA (<u>**HMG-CoA**</u>) reductase and reduce production of mevalonate-derived isoprenoids required for prenylation of multiple signaling proteins, including Rho-family GTPases[19, 20]. Through this cholesterol-independent mechanism, statins can alter Rho-family GTPase localization, cytoskeletal signaling, and endothelial inflammatory responses. Although statins have shown endothelial-protective effects in experimental systems[21-24], randomized clinical trials in unselected patients with established sepsis[25] or acute respiratory distress syndrome (<u>**ARDS**</u>)[26] have not demonstrated consistent clinical benefit, suggesting that statin-sensitive endothelial mechanisms may depend on cell type, timing, statin species, inflammatory phenotype, or drug delivery. In this study, we tested whether simvastatin protects tight-junction-forming HDMECs from TNF-induced barrier dysfunction and whether this protection involves mevalonate-dependent regulation of RhoB localization and activation.

## RESULTS

### Simvastatin attenuates TNF-induced barrier loss in HDMEC monolayers

To determine whether simvastatin protects microvascular endothelial barrier function during inflammatory stimulation, HDMEC monolayers were pretreated with simvastatin or vehicle for 24 hours and then stimulated with TNF for 6 hours while the pretreatment conditions were maintained. Barrier function was monitored in real time by TEER. TNF induced a progressive decline in normalized resistance, whereas simvastatin pretreatment attenuated this decrease (Figure 1A).

**Fig 1.**
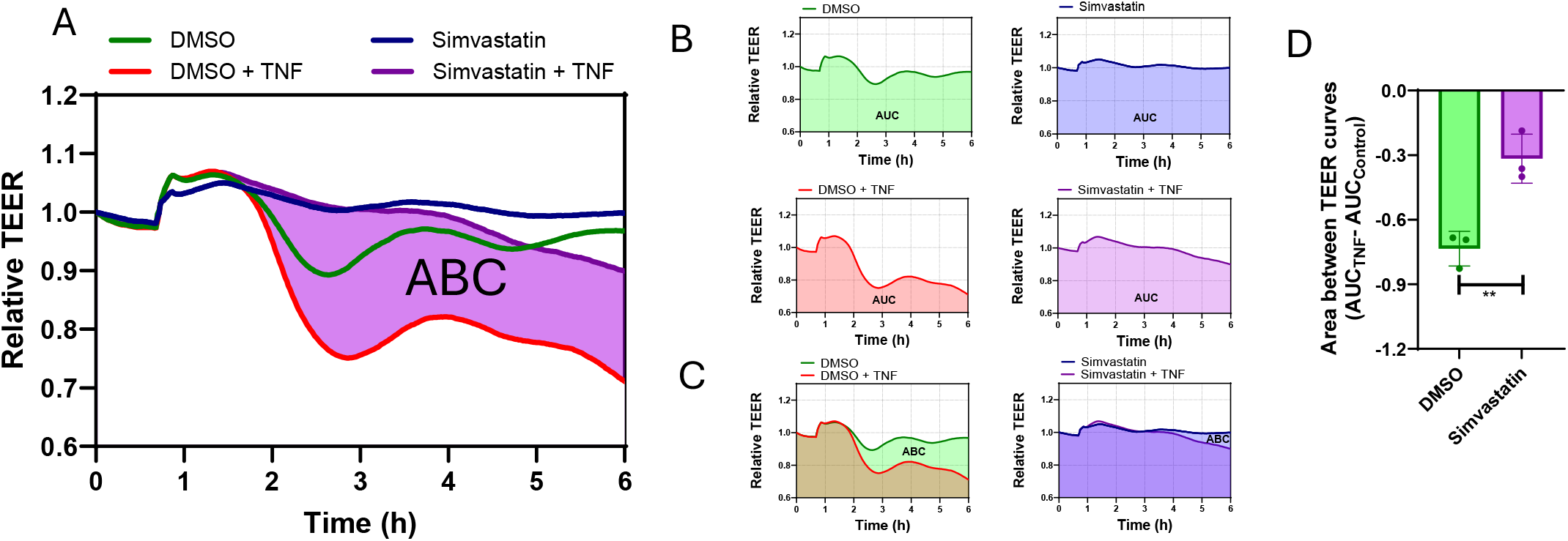
Simvastatin attenuates TNF-induced capillary leak. Relative TEER changes **(A)**, area under relative TEER curves **(B)**, area between relative TEER curves **(C)**, and quantification of area between relative TEER curves **(D)** of HDMECs pretreated ± simvastatin (1uM) for 24h followed by ± TNF stimulation (10ng/ml) for 6h. After simvastatin pretreatment, cells remained in the same medium and TNF was added directly to the wells. Representative of n= 3 (A, B, C) and n=3 (D). p = **≤0.01

Quantification of normalized AUC demonstrated that TNF reduced integrated barrier function over the 6 hour stimulation period, whereas simvastatin pretreatment preserved TEER (Figure 1B). The area between the TNF and simvastatin plus TNF curves further confirmed that simvastatin significantly reduced the magnitude of TNF-induced barrier loss (Figure 1C,D).

To confirm that simvastatin engaged the expected intracellular response, we measured LDL receptor expression and SREBP2 processing. Simvastatin increased LDL receptor expression and increased nuclear SREBP2 signal, consistent with inhibition of cholesterol/mevalonate pathway signaling (Figure S1). These control experiments support that the observed effects occurred under conditions in which simvastatin produced its expected cellular response.

### Simvastatin preserves tight junction organization and reduces RhoB membrane association

We next examined whether the TEER preservation observed with simvastatin was accompanied by preservation of tight junction organization. Under basal conditions, ZO-1 forms a continuous pattern along endothelial cell borders, consistent with intact intercellular junctions. TNF stimulation disrupted this smooth, continuous ZO-1 pattern and produced irregular, discontinuous junctional staining. Simvastatin pretreatment preserved more continuous ZO-1 staining after TNF stimulation, indicating attenuation of TNF-induced junctional disruption (Figure 2A). Similar preservation of junctional staining was observed for claudin-5 in supporting experiments (Figure S2).

**Fig 2.**
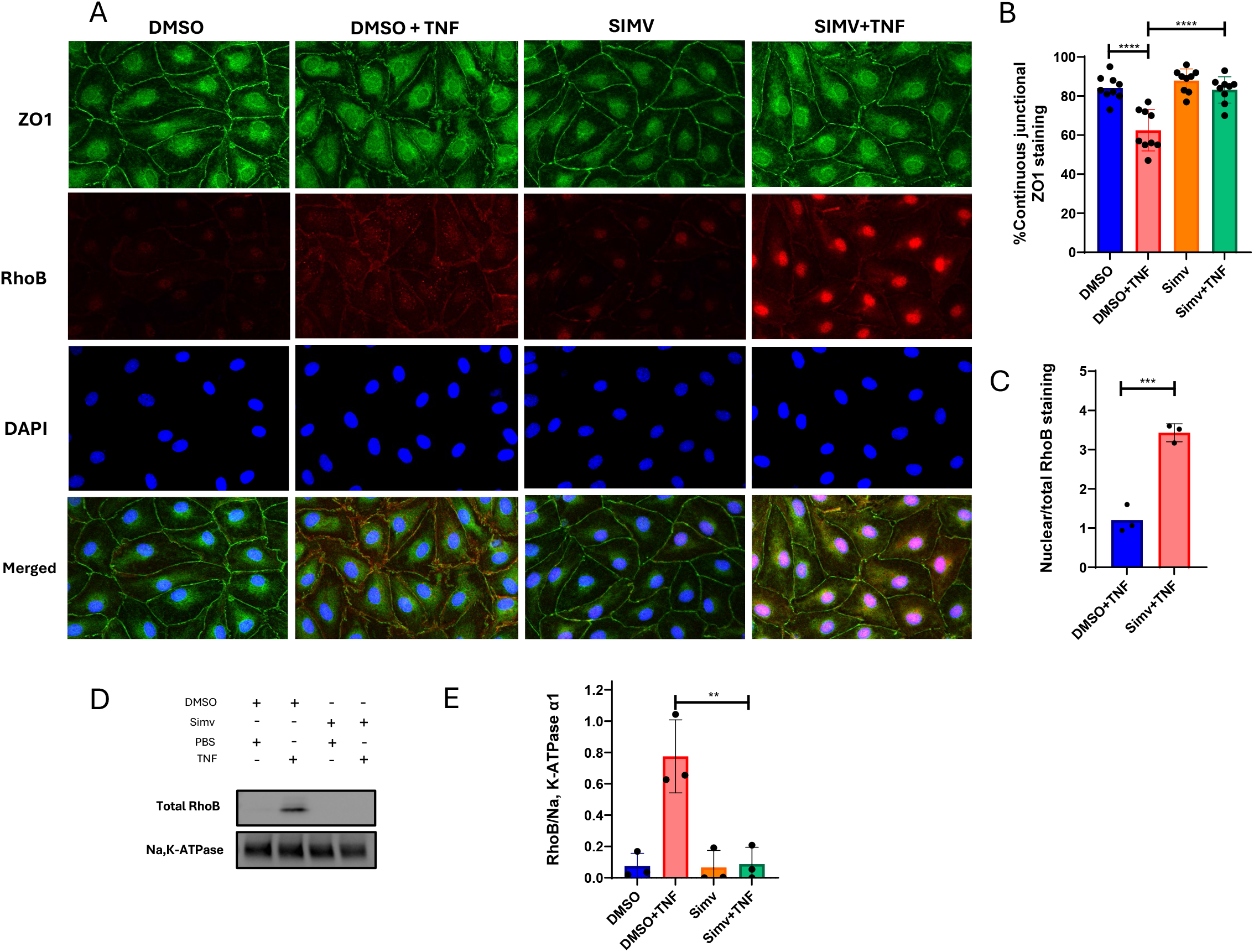
Simvastatin preserves tight junction organization and reduces RhoB membrane association. Confocal immunofluorescence staining of RhoB and ZO1 **(A)**, percentage of continuous junctional of ZO-1 staining **(B)**, nuclear vs non-nuclear staining of RhoB **(C)**, western blot of membrane-localized RhoB **(D)** and densitometric quantification of total RhoB protein expression normalized to Na,K-ATPase alpha1 (plasma membrane marker) **(E)** in HDMECs pretreated ± simvastatin (1uM) for 24h followed by ± TNF stimulation (10ng/mL) for 6h. Representative of n=3 (A,D) and n=3 (B,C,E). p= ** ≤ 0.01;*** ≤ 0.001; ****≤ 0.0001

RhoB staining also differed across treatment conditions. TNF increased RhoB signal at the cell periphery, whereas simvastatin pretreatment reduced peripheral RhoB staining and increased the relative nuclear or perinuclear RhoB signal (Figure 2A). Quantitative image analysis showed that simvastatin increased the proportion of cell borders with smooth, continuous ZO-1 staining after TNF stimulation (Figure 2B). Simvastatin also increased the nuclear-to-non-nuclear RhoB fluorescence ratio compared with TNF-stimulated cells (Figure 2C). Consistent with these findings, subcellular fractionation followed by Western blot analysis showed that simvastatin markedly reduced RhoB abundance in the membrane-enriched fraction, with Na^+^/K^+^-ATPase alpha1 used as a membrane marker and loading control (Figure 2D,E).

To test whether RhoB contributes to the same barrier-disruptive pathway targeted by simvastatin, we compared RhoB silencing with simvastatin treatment. RhoB knockdown attenuated TNF-induced TEER loss to a degree comparable to simvastatin pretreatment. Combining RhoB knockdown with simvastatin did not provide additional protection beyond either intervention alone (Figure S3D). These data support the interpretation that simvastatin protects HDMEC barrier function, at least in part, by modulating a RhoB-dependent pathway.

### Simvastatin suppresses TNF induced RhoB GTP loading and MLC phosphorylation

Because RhoB membrane association is required for efficient interaction with upstream regulators and downstream effectors, we next tested whether simvastatin altered RhoB activation. Total RhoB and GTP-bound RhoB were measured in HDMEC lysates after simvastatin pretreatment and TNF stimulation. TNF increased

GTP-bound RhoB, whereas simvastatin markedly reduced GTP-bound RhoB (Figure 3A,B). In contrast, total RhoB abundance was not significantly reduced under these conditions after normalization to loading control (Figure 3C,D). Thus, simvastatin decreased the active pool of RhoB without a corresponding significant decrease in total RhoB protein. We also examined phosphorylation of myosin light chain 2 at Thr18/Ser19, a downstream marker of actomyosin contractility associated with endothelial barrier disruption. TNF increased MLC phosphorylation, whereas simvastatin pretreatment markedly reduced this response (Figure 3C,E). These findings are consistent with suppression of RhoB-associated contractile signaling and with the preservation of endothelial barrier function observed by ECIS.

**Fig 3.**
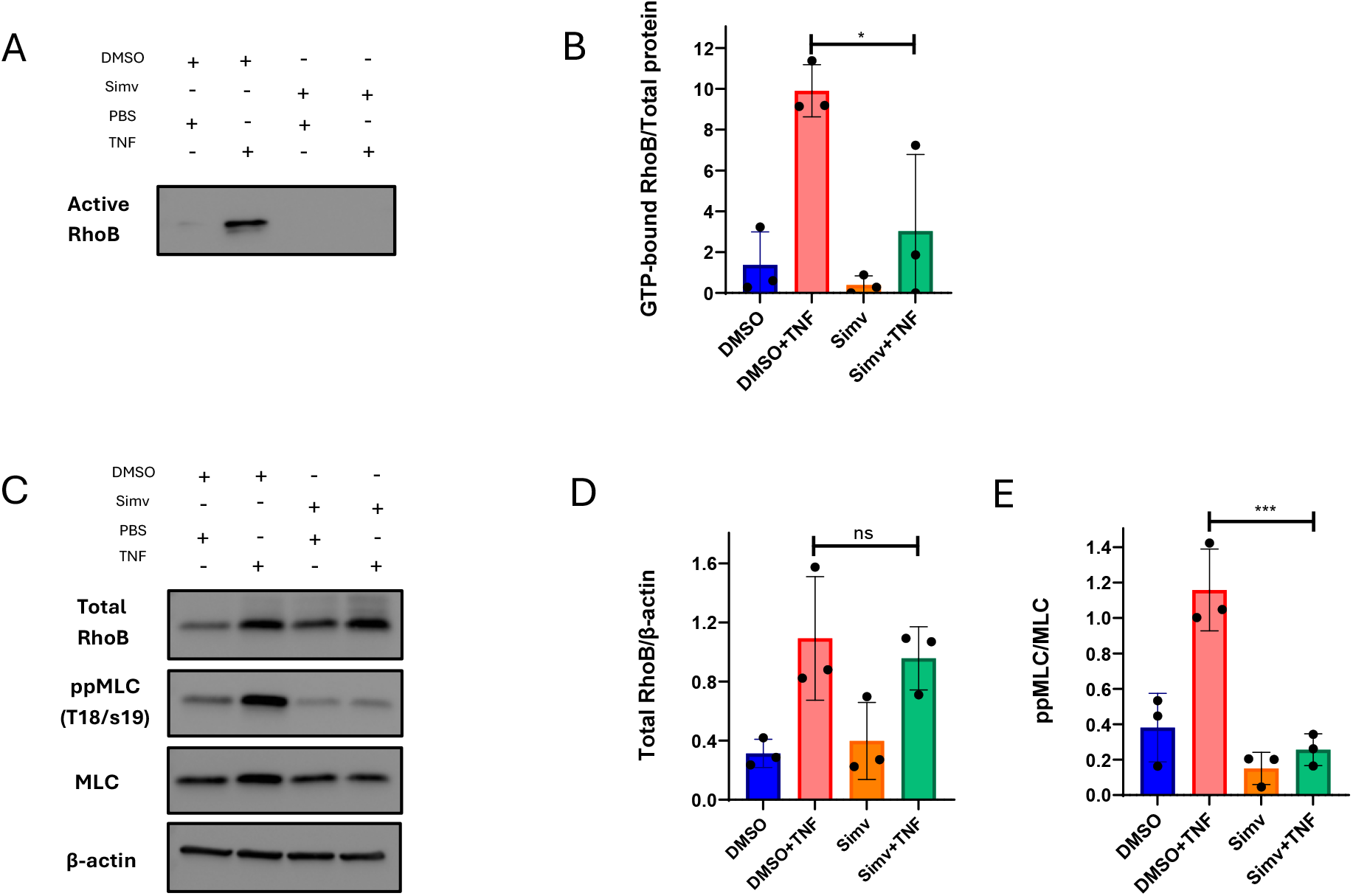
**Simvastatin reduces GTP-bound RhoB without a statistically detected reduction in total RhoB**, western blot of GTP-bound RhoB **(A)**, densitometric quantification of GTP-bound RhoB expression normalized to total protein **(B)**, western blot of total RhoB, ppMLC, MLC and β-actin **(C)**, densitometric quantification of total RhoB protein expression normalized to β-actin **(D)** and ppMLC protein expression normalized to total MLC **(E)** of HDMECs pretreated ± simvastatin (1uM) for 24h followed by ± TNF stimulation (10ng/ml) for 6h. Representative of n= 3 (A, C) and n= 3 (B, D, E). p = *≤0.05 ;*** ≤ 0.001.

### Mevalonate reverses simvastatin-mediated RhoB redistribution and barrier protection

To test whether simvastatin-mediated barrier protection required inhibition of the mevalonate pathway, HDMECs were supplemented with mevalonate during simvastatin pretreatment before TNF stimulation (Figure 4A). TNF reduced normalized TEER, whereas simvastatin pretreatment attenuated this loss. Mevalonate supplementation reversed the protective effect of simvastatin, resulting in TEER decline after TNF stimulation (Figure 4B). Quantification of AUC confirmed that mevalonate significantly reduced simvastatin-mediated preservation of barrier function (Figure 4C).

**Fig 4.**
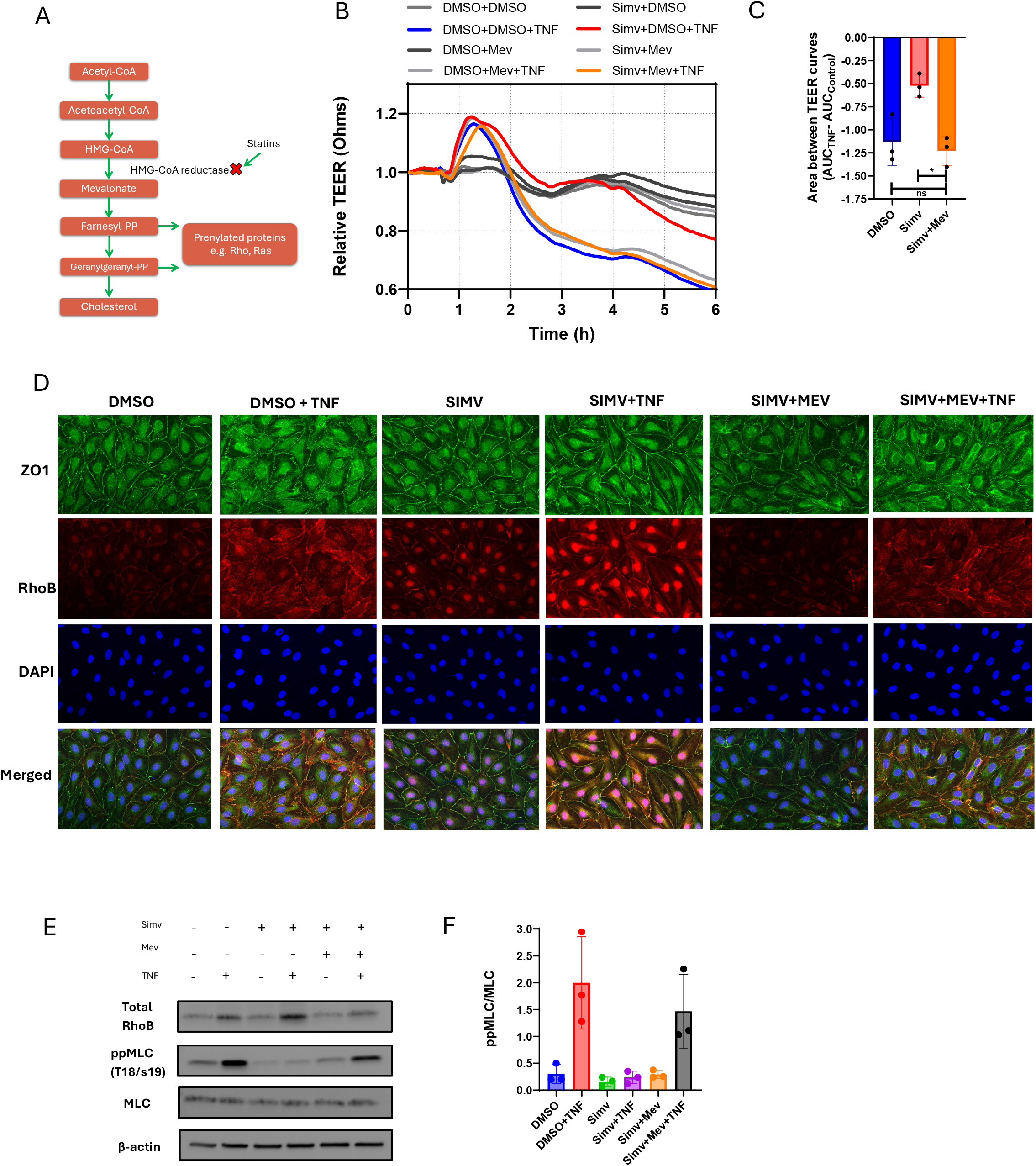
Mevalonate rescues TNF-induced capillary leak. Schematic diagram of mevalonate pathway and statin effect **(A)**. Relative TEER changes **(B)**, quantification of area between relative TEER curves **(C)**, Confocal immunofluorescence staining of ZO-1 and RhoB **(D)**, Western blot of RhoB, ppMLC, MLC and β-actin **(E)**, densitometric quantification of ppMLC normalized to total MLC **(F)** in HDMECs pretreated ± simvastatin (1uM) ± mevalonate (200uM) for 24h followed by ± TNF stimulation (10ng/ml) for 6h. Representative of n=3 (B, D, E) and n=3 (C, F, G). p = *≤0.05

We next examined whether mevalonate also reversed the effects of simvastatin on junctional organization and RhoB localization. TNF disrupted continuous ZO-1 staining at endothelial borders and increased peripheral RhoB signal. Simvastatin preserved ZO-1 continuity and reduced peripheral RhoB localization after TNF stimulation. Mevalonate supplementation reversed these effects, restoring peripheral RhoB signal and coinciding with renewed ZO-1 disruption (Figure 4D).

Consistent with these imaging findings, mevalonate supplementation restored TNF-induced MLC phosphorylation in simvastatin-treated cells (Figure 4E,F). Together, these findings indicate that restoration of the mevalonate pathway reverses the effects of simvastatin on RhoB localization, MLC phosphorylation, and endothelial barrier function. These data support a model in which mevalonate-dependent lipid modification of signaling proteins, including Rho-family GTPases, contributes to simvastatin-mediated barrier protection.

## DISCUSSION

In the present study, we show that simvastatin preserves human microvascular endothelial barrier function during TNF stimulation. Simvastatin attenuated TNF-induced TEER loss, preserved TJ organization, reduced membrane-associated and GTP-bound RhoB, and suppressed MLC phosphorylation. These protective effects were abolished by mevalonate supplementation. Together, these findings suggest that simvastatin protects TJ-forming microvascular endothelial monolayers by disrupting mevalonate-dependent RhoB membrane localization and activation.

RhoB differs from closely related Rho GTPases in its dynamic expression, lipid modification, and intracellular distribution. Whereas RhoA and Cdc42 are most often described as cytosolic proteins that undergo stimulus-dependent recruitment to membranes[27, 28], RhoB localizes to multiple membrane compartments, including the plasma membrane[28-30], Golgi[28], early and late endosomes and multivesicular bodies[29, 31-33]. Nuclear RhoB has also been reported[34, 35]. This compartmentalization is biologically important because GTP-loaded RhoB has been detected at both plasma membrane and endosomal membranes, and RhoB can signal from distinct subcellular compartments[30, 36, 37]. These observations support the concept that RhoB signaling is controlled not only by GTP loading, but also by localization.

RhoB membrane association is regulated in part by post-translational modification of its C-terminal CAAX motif. Unlike many Rho-family GTPases, RhoB can undergo either farnesylation or geranylgeranylation, modifications that influence its membrane targeting and signaling properties[32, 38-40]. Fluorescence loss in photobleaching and live-cell imaging studies further suggest dynamic exchange between plasma membrane and endosomal RhoB pools, with plasma membrane-associated RhoB supplied in part by intracellular endosomal compartments[36]. Thus, RhoB provides a useful model for testing whether disruption of membrane targeting can alter inflammatory endothelial barrier signaling.

Our findings extend this spatial model by showing that simvastatin alters RhoB localization during TNF stimulation. TNF increased peripheral RhoB staining and membrane-associated RhoB, whereas simvastatin reduced peripheral and membrane-associated RhoB and increased the relative nuclear or perinuclear RhoB signal. Because simvastatin inhibits the mevalonate pathway, these findings are consistent with altered isoprenoid-dependent membrane targeting of RhoB or proteins regulating RhoB trafficking. We did not directly measure RhoB prenylation, however, and therefore cannot distinguish altered RhoB lipid modification from effects on trafficking, binding partners, or membrane organization. These mechanisms are not mutually exclusive.

The increase in nuclear or perinuclear RhoB signal is intriguing. Prior studies have reported nuclear RhoB localization, and nuclear RhoB-GTP has been implicated in endothelial transcriptional regulation through VEZF1[35, 36]. Nevertheless, the functional significance of nuclear or perinuclear RhoB in our system is unclear; future studies examining the prenylation state, activation state, mobility, and binding partners of nuclear-associated RhoB will be required to determine the functional significant of nuclear RhoB.

Despite these changes in localization, simvastatin did not significantly affect total RhoB protein abundance under the conditions tested. Instead, simvastatin markedly reduced GTP-bound RhoB. These findings suggest that the reduction in recovered GTP-bound RhoB cannot be explained solely by a reduction in total RhoB abundance. Together with the reduction in membrane-associated RhoB, the data are consistent with mevalonate pathway inhibition limiting RhoB activation at membrane-associated signaling compartments.

The functional consequences of altered RhoB localization and activation were reflected in preservation of endothelial barrier function. By reducing membrane-associated, GTP-bound RhoB, simvastatin may limit engagement of downstream contractile effectors that destabilize tight junctions. Consistent with this interpretation, simvastatin attenuated TNF-induced MLC phosphorylation, preserved ZO-1 and claudin-5 organization, and reduced TEER loss. These findings extend our prior work demonstrating that TNF-induced ArhGEF10 selectively activates RhoB to disrupt tight junctions and impair barrier function in HDMECs[11].

Mevalonate supplementation provided further support for involvement of the mevalonate pathway. Restoration of mevalonate availability reversed simvastatin-mediated TEER preservation and was accompanied by restored peripheral RhoB localization, renewed ZO-1 disruption, and restored MLC phosphorylation. These findings strengthen the association between mevalonate pathway inhibition, altered RhoB signaling, and preservation of endothelial barrier integrity. However, mevalonate rescue does not identify the responsible isoprenoid species or protein substrate. Direct measurement of RhoB lipid modification and branch-specific rescue with FPP or GGPP will be required to determine whether altered RhoB prenylation mediates these effects.

Beyond its effects on cytoskeletal signaling, RhoB also regulates endothelial inflammatory responses. RhoB is induced by inflammatory stimuli, and RhoB depletion attenuates TNF-induced p38 activation, adhesion molecule expression, and inflammatory cytokine production[10, 41]. Thus, statin-sensitive regulation of RhoB may influence both the mechanical and inflammatory components of endothelial dysfunction, although the present study directly examined only barrier-associated signaling.

The present findings may be relevant to inflammatory critical illness, although they do not establish a therapeutic effect of simvastatin in patients. Secondary analyses of ARDS trials have identified hyperinflammatory phenotypes characterized by systemic inflammation, endothelial activation, shock, and greater organ dysfunction[42-44], raising the possibility that responses to vascular-directed therapies differ across biological phenotypes. In HARP-2, a secondary analysis suggested greater benefit from simvastatin within a hyperinflammatory subgroup[45]; however, these findings were hypothesis-generating and did not establish a predictive treatment phenotype. More broadly, randomized trials have not demonstrated consistent clinical benefit from initiating statins in unselected patients with established ARDS. Rosuvastatin did not improve clinical outcomes in sepsis-associated ARDS in SAILS, and simvastatin did not improve the primary outcome in unselected ARDS in HARP-2[25, 26]. These clinical results contrast with extensive experimental evidence that statins alter endothelial inflammatory signaling, cytoskeletal regulation, and vascular permeability[46-52]. One explanation is that statin-sensitive endothelial mechanisms are context dependent and influenced by drug, vascular bed, inflammatory state, treatment timing, and tissue exposure. Our findings identify mevalonate-dependent regulation of RhoB localization and activation as one such mechanism in human microvascular ECs.

Several limitations constrain interpretation of these findings. First, HDMECs provide a tractable model of TJ-forming human microvascular endothelium[8], but endothelial phenotypes vary substantially among vascular beds[53-55], and these findings cannot be assumed to apply to pulmonary or other organ-specific endothelium. Second, the micromolar simvastatin concentration used *in vitro* exceeds circulating concentrations achieved during conventional systemic therapy[56], limiting direct pharmacological extrapolation. Third, simvastatin was administered before TNF stimulation, establishing protection against developing barrier injury rather than rescue of established dysfunction. Finally, we did not directly measure RhoB prenylation. Mevalonate rescue together with altered RhoB membrane association and activation supports involvement of the mevalonate pathway but does not establish RhoB lipid modification as the responsible molecular intermediate.

In conclusion, simvastatin preserves TJ-forming EC barrier function during TNF stimulation while reducing membrane-associated and GTP-bound RhoB and downstream MLC phosphorylation. Mevalonate reverses these effects, linking endothelial barrier protection to the mevalonate pathway. Together, these findings identify spatial regulation of RhoB as a statin-sensitive mechanism associated with inflammatory microvascular barrier dysfunction and provide a mechanistic framework for investigating RhoB-dependent vascular injury during inflammatory critical illness.

## MATERIALS AND METHODS

### Cells, treatments, and transfection

Primary human dermal microvascular endothelial cells (<u>**HDMECs**</u>) were isolated from de-identified normal adult skin obtained through the Yale University Department of Pathology Tissue Collection Service using established protocols[8]. The Yale University Institutional Review Board classified this work as non-human subjects research. HDMECs were cultured in EBM-2 Endothelial Cell Growth Basal Medium (Lonza, CC-3156) supplemented with EGM-2 MV Microvascular Endothelial Cell Growth Medium-2 SingleQuots (Lonza, CC-4147) and used between passages 3 to 8. Where indicated, cells were treated with recombinant human TNF (Gibco; 10 ng/mL), simvastatin - 2,2-dimethyl-1S,2,3R,7S,8S,8aR-hexahydro-3,7-dimethyl-8-[2-[(2R,4R)-tetrahydro-4-hydroxy-6-oxo-2H-pyran-2-yl]ethyl]-1-naphthalenyl ester, butanoic acid- (Cayman Chemical; 1 µM), mevalonate (Cayman Chemical; 200 µM), or vehicle control. Simvastatin and mevalonate were added 24 h before TNF stimulation and maintained during TNF treatment unless otherwise indicated. Vehicle concentration was matched across treatment groups and did not exceed [0.01%]. For knockdown experiments, HDMECs were transfected with non-targeting or RhoB-targeting SMARTpool siRNA (Dharmacon; 40 nM) using Lipofectamine RNAiMAX (Invitrogen) according to the manufacturer’s instructions. Cells were analyzed 48 h after transfection.

### Transendothelial electrical resistance measurement

Transendothelial electrical resistance (<u>**TEER**</u>) of HDMEC monolayers was measured using electric cell-substrate impedance sensing (<u>**ECIS**</u>; Applied BioPhysics). HDMECs were seeded at 3 × 10^4^ cells per well onto gelatin-coated 96-well gold electrode arrays (Applied BioPhysics, 96W20idfPET). TEER was monitored daily until resistance reached a stable plateau, indicating formation of a mature endothelial barrier. Treatments were added without replacing the medium during the preceding 24 h. Resistance was measured every 60 s using a 1 µA constant alternating current at 4000 Hz with an ECIS Z-Theta instrument and ECIS software version 1.2.243 0PC (Applied BioPhysics). TEER values were normalized to the resistance immediately before TNF addition, which was assigned a value of 1.0. For each experiment, technical replicate wells were averaged after normalization. Area under the curve (<u>**AUC**</u>) and area between curves were calculated from normalized TEER traces over the indicated time interval using GraphPad Prism. Area between curves (<u>**ABC**</u>) was calculated as the integrated difference between treatment and control AUC over the time period described. ABC analysis allows for comparison of effect over time relative to control condition across unique biologic replicates. Normalization enabled comparison across HDMEC donors and ECIS arrays.

### Membrane fractionation

Membrane-associated RhoB was assessed using Subcellular Protein Fractionation Kit for Cultured Cells (Thermo Scientific™”). Briefly, HDMECs were harvested with trypsin, centrifuged at 500 × g for 5 min, and washed with ice-cold PBS. Cell pellets were resuspended in cytoplasmic extraction buffer (CEB) and incubated at 4°C for 10 min with gentle mixing, followed by centrifugation at 500 × *g* for 5 min. The supernatant containing the cytoplasmic fraction was removed, and the remaining pellet was resuspended in ice-cold membrane extraction buffer (MEB) supplemented with protease inhibitors. Samples were vortexed for 5 s and incubated at 4°C for 10 min with gentle mixing, followed by centrifugation at 3,000 × *g* for 5 min. The resulting supernatant, corresponding to the membrane-enriched fraction, was collected and subjected to immunoblot analysis for RhoB. Na^+^/K^+^-ATPase α1 was used as a membrane loading control.

### Western blotting (WB)

HDMECs were lysed directly in Laemmli sample buffer containing β-mercaptoethanol. Lysates were sonicated, boiled at 95°C for 5 min, and separated by SDS-PAGE using Bio-Rad precast gels. Proteins were transferred to polyvinylidene difluoride membranes (PVDF; Bio-Rad). Membranes were blocked in 5% bovine serum albumin (BSA) in Tris-buffered saline with 0.1% Tween-20 (TBST) for 1 h at room temperature and incubated overnight at 4°C with primary antibodies against RhoB (1:500; Cell Signaling Technology; catalog no. 63876), phospho-myosin light chain 2 Thr18/Ser19 (1:1000; Cell Signaling Technology; catalog no. 95777), myosin light chain 2 (1:500; Cell Signaling Technology; catalog no. 8505), SREBP2 (1:500; Cell Signaling Technology; catalog no. 25940), Na,K-ATPase alpha1(1:500; Cell Signaling Technology; catalog no. 23565) and β-actin (1:50,000; Sigma Life Sciences; catalog no. A2228). After washing, membranes were incubated for 1 h at room temperature with HRP-conjugated anti-rabbit or anti-mouse secondary antibodies (1:1000; Cell Signaling Technology). Immunoreactive bands were detected using SuperSignal West Femto or SuperSignal West Pico PLUS chemiluminescent substrate (Thermo Fisher Scientific) and visualized using G:BOX imaging system and GeneSys software (Syngene). Densitometry was performed using ImageJ, and target proteins were normalized to β-actin, total protein concentration, or Na,K-ATPase alpha1 as appropriate.

### Quantitative polymerase chain reaction (qPCR)

Total RNA was extracted using the RNeasy Plus kit (Qiagen), and cDNA was synthesized using the High-Capacity cDNA Reverse Transcription Kit (Applied Biosystems). qPCR was performed using TaqMan probes for LDLR and 18S rRNA on a CFX Connect Real-Time PCR Detection System (Bio-Rad). Relative expression was calculated using the ΔΔCt method with 18S rRNA as the reference gene.

### Confocal Immunofluorescence microscopy

HDMEC monolayers were cultured on fibronectin-coated glass coverslips and fixed with 4% paraformaldehyde for 10 min at 37°C. Cells were washed three times with phosphate-buffered saline (PBS), permeabilized with 0.1% Triton X-100 in PBS for 10 min, washed again, and blocked with 2.5% BSA for 1 h at room temperature. Cells were incubated overnight at 4°C with primary antibodies against RhoB (1:100; Cell Signaling Technology; catalog no. 63876), Alexa Fluor 488-conjugated ZO-1 (1:50; Invitrogen; catalog no. 339188), or claudin-5 (1:100; Cell Signaling Technology; catalog no. 49564). After washing, cells were incubated for 1 h at room temperature with Alexa Fluor 568- or Alexa Fluor 647-conjugated secondary antibodies (Invitrogen). Coverslips were mounted using ProLong antifade mounting medium containing DAPI (Invitrogen). Images were acquired using a Leica TCS SP5 laser-scanning confocal microscope with a 20× objective. Laser settings, gain, offset, and exposure parameters were held constant across treatment groups within each experiment. ZO-1 continuity was quantified from randomly selected fields by measuring the proportion of cell borders with smooth, continuous junctional staining, as previously described. RhoB localization was quantified by segmenting DAPI-positive nuclear regions and measuring RhoB fluorescence intensity within nuclear and non-nuclear compartments using ImageJ. Image analysis was performed not blinded to treatment group.

### Pull-down assays of GTP-bound (active) GTPases

GTP-bound RhoB was measured using a Rho activation assay kit (Cytoskeleton, RT02-A) according to the manufacturer’s protocol. Briefly, HDMECs cultured in 6-well plates were lysed in the kit lysis buffer supplemented with protease inhibitors. Lysates were clarified by centrifugation, and an aliquot was reserved to determine total protein concentration. Equal volumes of each sample were incubated with Rhotekin-RBD agarose beads for 1 hour at 4°C with gentle agitation. Beads were pelleted by centrifugation and washed three times with kit wash buffer. Bound proteins were eluted in 30 µL reducing Laemmli sample buffer, heated at 95°C for 2 minutes, and resolved by SDS-PAGE. GTP-bound RhoB was detected by immunoblotting. In parallel, HDMECs cultured in 6-well plates were lysed with Laemmli sample buffer containing β-mercaptoethanol as a total input.

## Statistical analysis

Statistical analyses were performed using GraphPad Prism version 10.3.0. Data are presented as mean ± SD unless otherwise indicated. Biological replicates represent independent HDMEC donors. Technical replicates were averaged within each biological replicate before statistical testing. Comparisons between two groups were performed using two-tailed unpaired or paired Student’s t tests, as appropriate. Comparisons among more than two groups were performed using one-way ANOVA followed by Tukey’s multiple-comparisons test. For ECIS experiments, statistical comparisons were performed on AUC or area-between-curves values calculated from normalized TEER traces over the indicated time interval. Normality was assessed using visual inspection. A p value < 0.05 was considered statistically significant.

## Abbreviations

ARDS: acute respiratory distress syndrome
EC: endothelial cell
ECIS: electric cell-substrate impedance sensing
FPP: farnesyl pyrophosphate
GGPP: geranylgeranyl pyrophosphate
HDMEC: human dermal microvascular endothelial cell
MLC: myosin light chain
TEER: transendothelial electrical resistance
TJ: tight junction
TNF: tumor necrosis factor
ZO-1: zonula occludens-1

