## Supplemental figures for "Simvastatin Preserves Human Microvascular Endothelial Barrier Function Through Mevalonate-Dependent Regulation of RhoB Localization and Activation"

**Fig S1. Simvastatin increases LDL receptor expression and activates sterol regulatory element-binding protein 2 (SREBP2).** Relative gene expression of LDL receptor measured by RT-qPCR and fold change calculated by $2^{-\Delta\Delta CT}$ method (**A**), western blot for SREBP2 (**B**) and densitometric quantification of nuclear SREBP2 protein expression normalized to β-actin of HDMECs ± simvastatin (1μM, 30 horas). n=3 (A, C), representative of n=3 (B). p = \*≤0.05; \*\* ≤ 0.01

**Fig S2. Simvastatin preserves claudin-5 integrity following TNF stimulation.** Confocal immunofluorescence staining of claudin-5 in HDMECs pretreated ± simvastatin (1μM) for 24h followed by ± TNF stimulation (10ng/ml) for 6h. Representative of n=3.

**Fig S3. RhoB silencing phenocopies simvastatin-mediated endothelial barrier protection.** Confocal immunofluorescence staining (**A**) and western blot of RhoB and β-actin (**B**) and densitometric quantification of total RhoB protein expression normalized to β-actin (**C**) of HDMECs treated ± RhoB siRNA ± TNF stimulation (10ng/ml) for 6h. Relative TEER changes (**D**) of HDMECs treated ± RhoB siRNA followed by ± simvastatin (1μM) and subsequent TNF stimulation (10ng/ml) for 6h. Representative of n=3 (A, B, D), n=3 (C), p = \*\*\* ≤ 0.001.

Figure S1

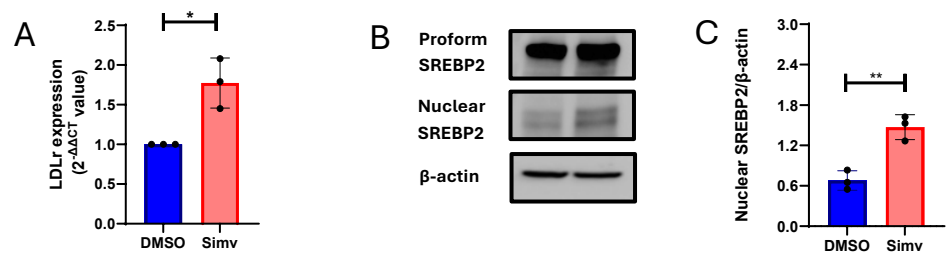

Figure S2

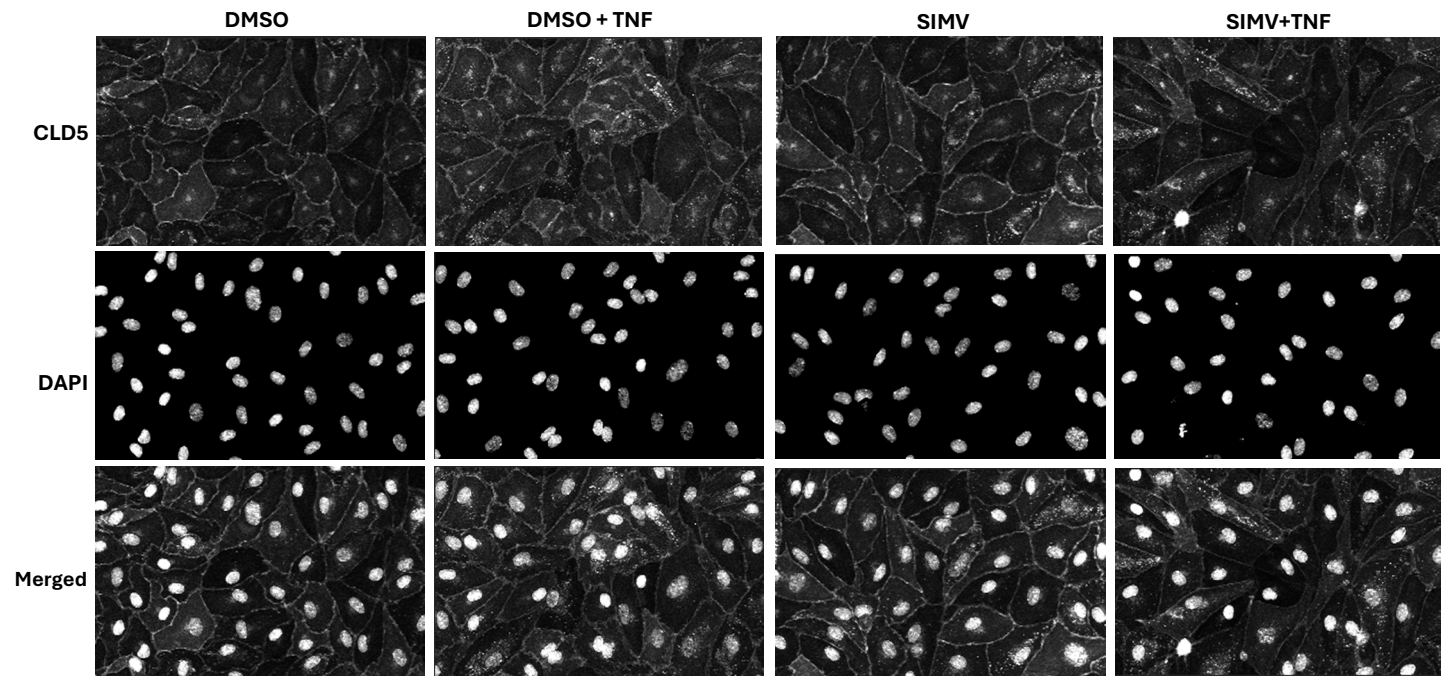

Figure S3

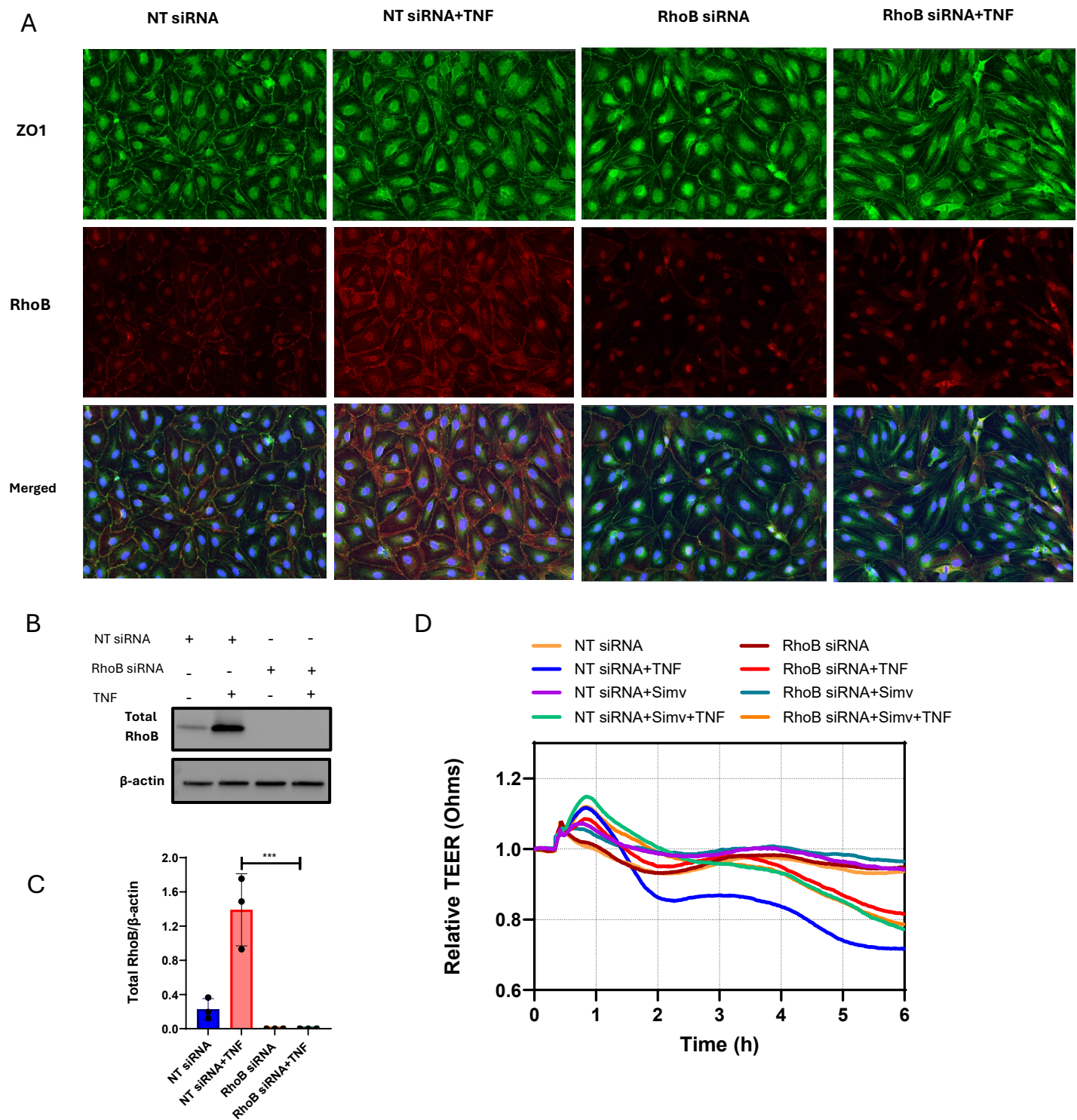
